# A shift between r and K strategies in the life cycle of a unicellular microalga

**DOI:** 10.1101/2024.09.24.614695

**Authors:** Lu Liu, Si Tang, Yaqing Liu, Boya Zhang, Yanlin Zhong, Dai Liu, Jin Zhou, Zhonghua Cai

**Affiliations:** Shenzhen Public Platform for Screening and Application of Marine Microbial Resources, Shenzhen International Graduate School, Tsinghua University, Shenzhen, 518055, Guangdong Province, PR China

## Abstract

r (high reproductivity but stress-sensitive offspring) and K (low reproductivity but high stress-resistant offspring) strategies are two classical life history theories. Contrary to the belief that one species only has a single strategy (i.e., either r or K), we present experimental evidence for the coexistence of both strategies in the isogenic microalga *Haematococcus lacustris*. Under standard conditions, swimming vegetative cells (SVC) normally grew and divided via binary fission (r strategy) and gradually shifted into non-swimming cysts (NSC) over time. Intriguingly, unlike the prevailing notion that NSC cannot propagate under stress, they were found to reproduce via multiple fission at a barely detectable growth rate, resulting in stress-resistant, non-mobile daughter cells and demonstrating a complete life history (K strategy). Collectively, our findings indicate that *H. lacustris* adopts both r and K strategies throughout its life history. This research enhances the understanding of the adaptation and survival of microbial populations in changing environments.

## Introduction

In order to survive and reproduce, microbial populations must both grow and defend, the switch between growth (reproduction) and stress resistance is therefore crucial to population fitness and has vital ecological consequences ^1,2^. Today, the changing environments, primarily driven by anthropogenic activities ^3,4^, pose unforeseen hurdles to living organisms. Understanding whether and how organisms adapt to such challenging environments, i.e., how microbial populations balance growth and defence, is a fundamental goal in ecology.

r and K theories are two classical reproduction frameworks describing distinct life history strategies ^5–7^, widely applied in microbiology, plant and animal research ^7–11^. In general, they describe a game between growth and stress resistance within the context of the battle for resources^12^. In detail, r-strategists are copiotrophs with the reproduction of more offspring but have lower individual-level stress resistance, an advantageous strategy in resource-replete environments ^11^. In contrast, K-strategists are oligotrophs with higher stress resistance at the expense of lower reproduction, an advantageous strategy in resource-limited environments ^13,14^. Empirically, most species are described to own only one life history strategy ^9,11,12,14,15^, i.e., either r or K strategy. However, from an evolutionary perspective, it would be theoretically logical to anticipate that populations would thrive and prosper more robustly if they were able to incorporate the advantages of r and K strategies in their life histories. In other words, it would be highly advantageous for a population to employ the r strategy under favourable conditions and shift to the K strategy under unfavourable ones, ultimately leading to an overall enhancement in population fitness.

Although such a presumption is promising, the discussion needs to be further delved into the study of individual species. Nonetheless, it seems possible for such a shift between r and K strategies in microbial populations. Specifically, many microbes are known for their relatively fast growth under favourable conditions (usually considered as r strategists) ^8,16^, whereas they can transform into a stress-resistant resting status under adverse conditions, e.g., spore formation ^17^. Here, the resting cell partially meet the definition of K strategy (high stress resistance). Nevertheless, since they are conventionally considered incapable of reproduction under adverse conditions ^18,19^, they cannot be fully classified as possessing a complete life history or as K strategists. Therefore, the shift between r and K strategies in one single microbial species requires more experimental evidence.

In this study, the team used an isogenic microalga *Haematococcus lacustris* (*H. lacustris*) as a model organism to explore the shift between r and K strategies. *H. lacustris* is a widely distributed, unicellular biflagellate freshwater microalga, and it is known for the ability to transform from growth-devoting swimming vegetative cells (SVC) into stress-resistant non-swimming cysts (a generalised term for unspecified dormant cells, NSC) ^20–22^, suitable for the goal of this study.

By integrating physiological, microscopic and transcriptomic data, here the team provide evidence that NSC were capable of reproduction under unfavourable conditions and showed a complete life history. Further evidence shows a reproduction mode shift between SVC and NSC, i.e., binary fission for SVC and multiple fission for NSC. Since the life histories of SVC and NSC exhibit a general congruence with the definition of both the r-strategy and K-strategy, respectively, it can thus be concluded that the shift between r and K strategies exists in this microalgal population. These findings challenge the prevailing view that stress-resistant NSC are incapable of reproduction under stressful conditions, and provide evidence in support of the shift between r and K strategies in the same microalgal strain, offering novel insights into the adaptation and maintenance of microbial populations.

## Results

### The switch from growth-devoting SVC to stress-resistant NSC

In general, consistent with previous reports ^9,11,12,14,15^, there was a phenotypic shift between *H. lacustris* populations over time grown under standard culture conditions, i.e., switching from SVC to NSC. Specifically, starting with nearly 100% of SVC (pyriform cells that were evenly distributed and showed a homogenous light green colour, Fig. S1A and S1C), the proportion of SVC decreased gradually with a substantial ratio of cells transformed into NSC (Fig. 1A, from day 7 to day 23). At the end of the experiment (day 23), almost all cells lost mobility (cells transformed into a spherical shape and precipitated to the bottom of the culture flask and formed an opaque layer of dark green cells, Fig. S1B and S1D). In addition, apart from the difference in mobility, cellular size, which have significant impacts on cellular functions ^23^, also showed drastic differences between these two phenotypes. In detail, results showed that NSC increased in size over time, as evidenced by the fact that cells larger than 25 µm in diameter only accounted for 8% of the population on day 1 but increased to 36% on day 23 (*t* test, *p* < 0.01). Simultaneously, in accordance with the increase in cellular size, the NSC exhibited a remarkably higher cellular weight, measuring 7.47 times higher than that of the SVC (Fig. 1C, t test, *p* < 0.01).

**Fig. 1.**
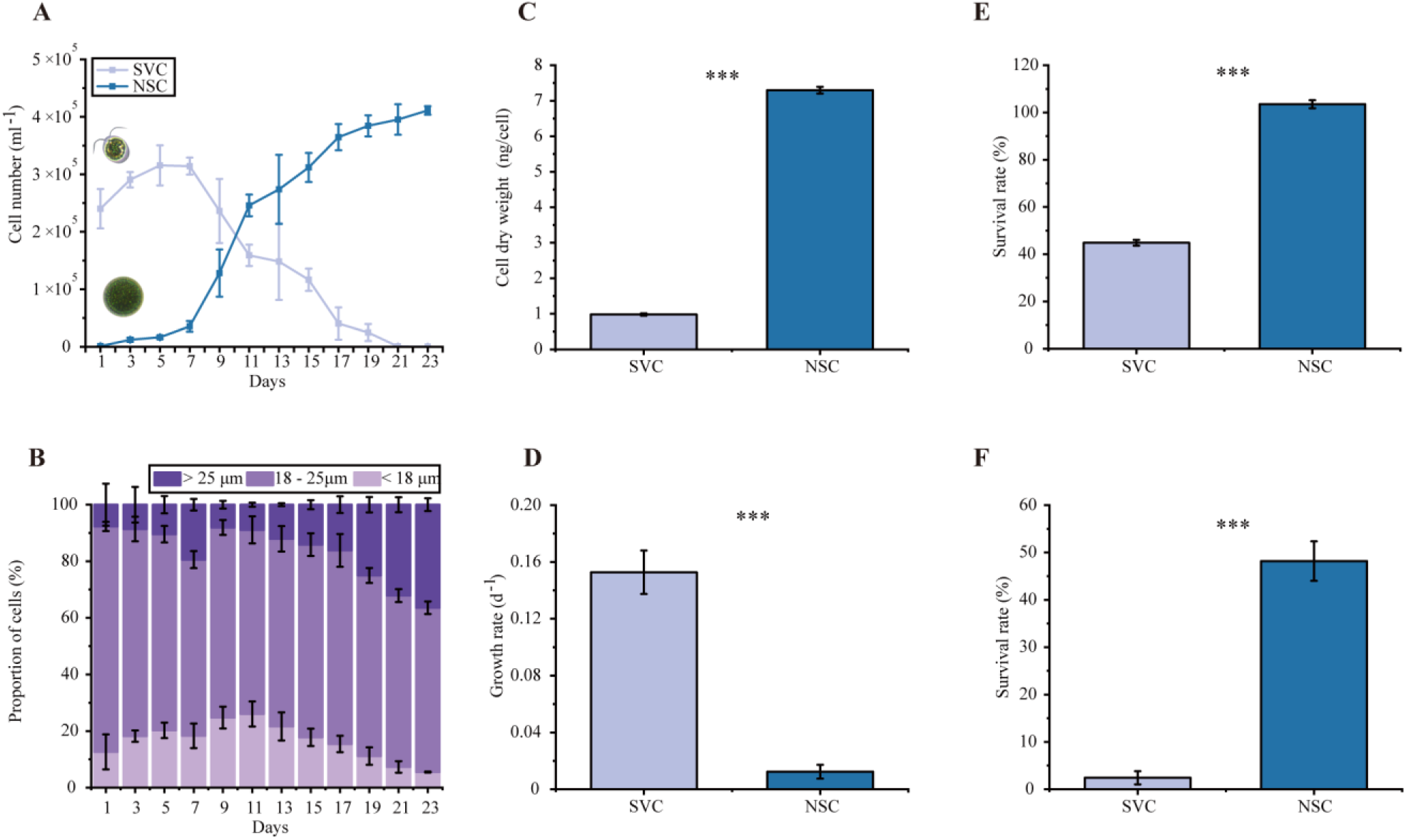
The growth-defence switch in *H. lacustris* populations. (A) Cell number dynamics of SVC (light blue) and NSC (dark blue) over a 23-day growth experiment. (B) Cellular size dynamics on every other day over a 23-day growth experiment (cell diameter greater than 25 μm, dark purple; cell diameter greater than 18 μm and smaller than 25 μm, violet; cell diameter smaller than 18 μm, lavender). (C) Cellular dry weight during a 6-day culture experiment. (D) Growth rate of each phenotype over a 6-day growth experiment. (E) Survival rate of each phenotype under high light irradiation. (F) Survival rate of each phenotype under drought stress. The error bars indicate the standard deviation of the triplicate values. N = 3 independent biological replicates. Statistical significance was calculated by Welch’s *t* test for pairwise comparisons of two treatments. Significance: * (*p* < 0.05), ** (*p* < 0.01), *** (*p* < 0.001).

To further explore the traits of each phenotype, we evaluated the growth performance of two phenotypes under either standard or stressful conditions, respectively. For growth rate, SVC can normally grow (0.15 d^-1^), while the increase in cell number of NSC was barely detectable under standard culture conditions (Fig. 1D). In contrast, under stressful conditions (high light irradiation and drought stress), NSC showed markedly higher survival rates than SVC. Specifically, after three days of exposure to high light irradiation, NSC had a survival rate of nearly 100%, whereas only 45% of the SVC remained viable (Fig. 2E, t test, *p* < 0.01). Here, dead SVC were bleached (Fig. S2A), whereas NSC retained intact cell shapes (Fig. S2B). Likewise, NSC showed significantly higher drought resistance (approximately 20-fold higher) than SVC (Fig. 1F, t test, *p* < 0.01).

**Fig. 2.**
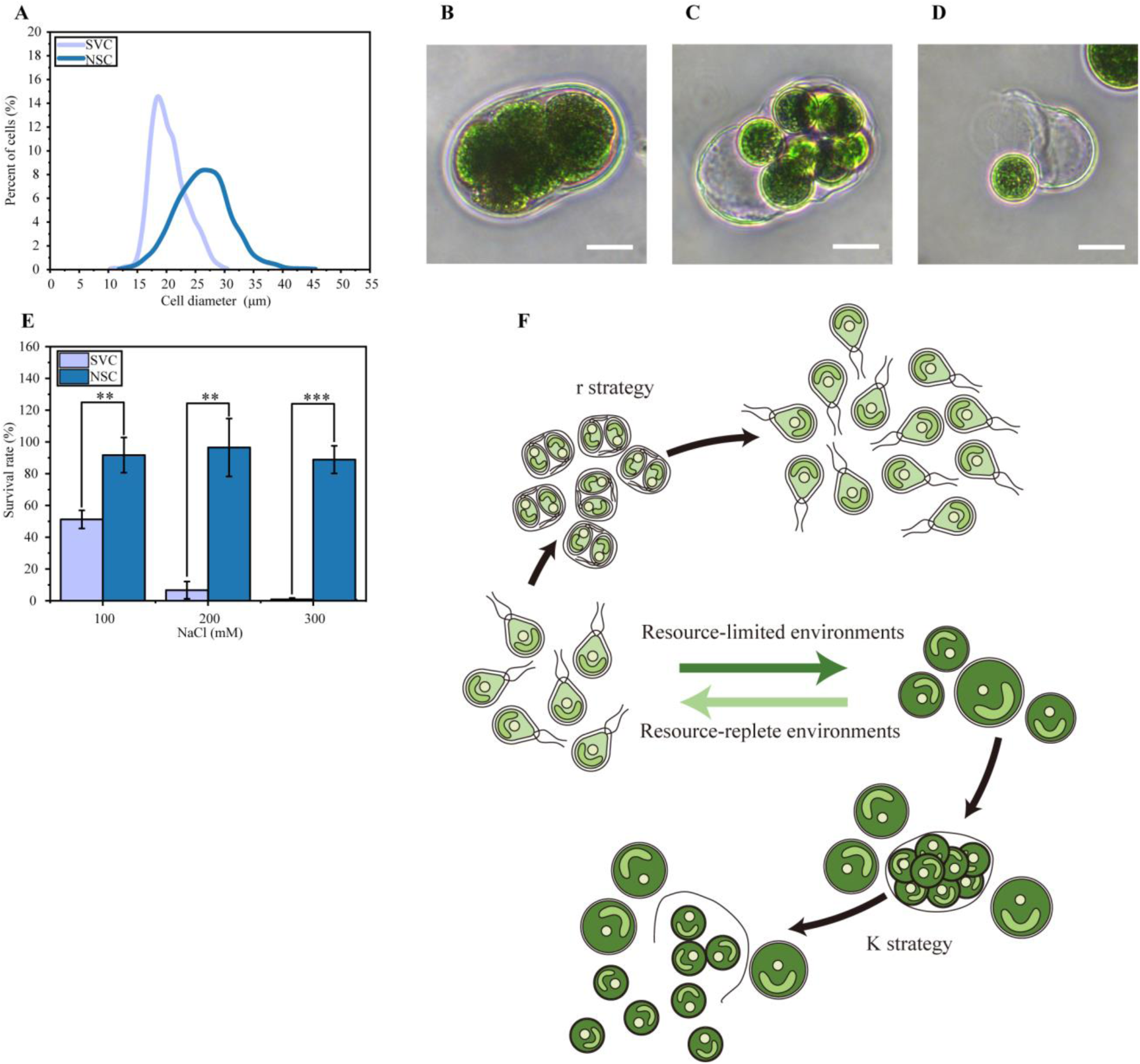
NSC reproduce via multiple fission. (**A**) The Gaussian kernel density curve illustrating the diameter distribution of SVC (light blue) and NSC (dark blue). (**B**) Microscopic images of NSC - propagation within the cyst. (**C**) Microscopic images of NSC - the preparation for offspring release. (**D**) Microscopic images of NSC - offspring release. (**E**) Survival rate of daughter cells from each phenotype under a NaCl gradient. (**F**) Schematic of the shift between r and K strategies with distinct reproduction modes. The error bars indicate the standard deviation of the triplicate values. N = 3 independent biological replicates. Statistical significance was calculated by Welch’s *t* test for pairwise comparisons of two treatments. Significance: * (*p* < 0.05), ** (*p* < 0.01), *** (*p* < 0.001). Scale bar: 20 μm.

Collectively, these results suggest that there was a growth and defence switch between the SVC and NSC. SVC are growth-devoted but sensitive to external stresses. Instead, NSC trade growth for stress resistance. Indeed, stress-resistant NSC were commonly regarded as resting cysts by other researchers ^18,24,25^, as such a phenotypical shift (from vegetative cells to resting cysts) is a ubiquitous strategy to cope with deteriorating environments in the microalgal phyla ^26^. From a population-level perspective, such a shift makes sense, because as time goes on, although no additional stresses are exerted on populations, their dwelling environments will deteriorate over time due to the increased stresses related to high cell density (such as reduced availability and increased competition for nutrients ^27^, self-shading ^28^, the release and accumulation of unfavourable substances ^29^) in older cultures, even initiated with favourable conditions. Notably, the traits of the SVC generally meet the characteristics of the r strategist, whereas NSC are not yet sufficient to be classified as K strategists, as they have been traditionally regarded as incapable of reproduction, thus cannot be defined to possess a complete life history strategy.

### NSC reproduce via multiple fission under stressful conditions

Traditionally, resting spores or cysts are believed to be incapable of reproduction under continuous stress, not only in the case of *H. lacustris* but also in other reported microalgal or bacterial species ^26,30–32^. In this study, we found that the increase in cell number of NSC was barely detectable (Fig. 1D), which seems to comply with the conventional idea. However, based on the results of cellular size dynamics (Fig. 1B), we were surprised by the fact that there was still a substantial proportion of NSC (25.2%, day 23) that were smaller than the average size of SVC (21.2 μm in diameter, day 1). This result is somehow counter-intuitive, as if NSC were solely derived from SVC, cells will gradually increase in size over time. As a consequence, all NSC should be generally larger than SVC and these observed small NSC should not exist. Accordingly, we speculated that these small cysts might derive from larger cysts, i.e., large NSC can somehow reproduce and release small daughter cysts. To test the presumption that NSC can reproduce, we first evaluated the cellular size distribution between SVC and older NSC (two months since all cells lost mobility). Theoretically, if all NSC were transformed from SVC, older NSC should be larger than SVC, as they have two months to grow bigger. In contrast, the existence of small cysts (not larger than the average diameter of SVC) would provide evidence that smaller NSC may derive from mother NSC but not from SVC, i.e., cysts can reproduce in the presence of stress. As shown in Fig. 2A, an overlapping cellular size distribution between the SVC and NSC was observed, indicating a substantial proportion of NSC were not bigger than SVC. Specifically, 9.9% of older NSC were smaller than the average diameter (20.4 μm) of SVC (Fig. 2A). In this case, the existence of small NSC cannot be explained by the prevailing idea that the cysts were only transformed from SVC ^33^. Instead, it suggests that NSC can reproduce and release daughter NSC on their own.

To gain more evidence for the reproduction ability of NSC under stress, the microscopic observation was conducted. Although not focused on the performance of the same mother cyst, we identified stepwise microscopic evidence showing how NSC reproduce and release daughter cysts: 1) propagation within the cyst (multiple daughter cells were produced within the mother cyst, Fig. 2B); 2) the preparation for the release of offspring (the outer layer of the mother cyst was to collapse and inner daughter cells were to be released, Fig. 2C); 3) the release of offspring (the outer layer of the mother cyst collapsed and daughter cysts were released, Fig. 2D). Notably, this reproduction mode, defined as multiple fission ^34,35^, is distinct from the binary fission of SVC. Furthermore, the mobility of offspring cells of each phenotype mirrored those of their respective mother cells. Specifically, offspring derived from SVC exhibited swimming capability, whereas offspring from NSC remained stationary (the whole population of the two-month-old culture was non-mobile). Importantly, it should be noted that the observed multiple fission of NSC under adverse conditions is totally distinct from cyst germination, as documented in previous studies ^36–38^. In cyst germination, the proliferation of swimming offspring through multiple fission occurs when NSC are transferred to more conducive environments, such as a fresh medium.

To further explore the traits of daughter NSC, we next evaluated the growth performance of daughter cells of NSC or SVC under stressful conditions (salinity). To this end, SVC at the exponential phase and NSC smaller than 21 µm in diameter (considered as newly released daughter cysts) were used for stress test. In general, similar to the performance of their mother cysts (Fig. 1E & 1F), daughter NSC showed a significantly higher stress resistance than daughter SVC (Fig. 2E, t test, *p* < 0.01). In particular, 48.8%, 93.3% and 99.1% of daughter SVC bleached or collapsed upon exposure to NaCl, whereas 91.7%, 96.4% and 88.9% of daughter NSC still remained intact in the presence of 100, 200, 300 mM of NaCl, respectively. These results suggest that although NSC propagate via multiple fission at a relatively lower rate compared to SVC, released daughter cysts are already prepared to survive under adverse conditions.

Finally, we investigated whether the observed NSC reproduction under stressful conditions is a strain-specific phenomenon. To achieve this, we included three additional *H. lacustris* strains (FACHB-797, FACHB-871, FACHB-1928) into our study and conducted a comparative analysis of cellular size distribution between SVC and NSC for each strain. As shown in Fig. S3, similar to the core strain (FACHB-712) used in this study, we found that NSC of all three additional strains also contained a substantial proportion of small cysts (37.6% for FACHB-797, 23.9% for FACHB-871 and 35.5% for FACHB-1928, respectively), which were smaller than the average size of their corresponding SVC. This result suggests that the reproduction of cysts under deteriorating conditions is not strain-specific but a general strategy of *H. lacustris*.

In summary, by integrating the results from cellular size comparison and direct microscopic observation, unlike the prevailing view that microalgal cysts cannot reproduce under adverse environments ^18,19^, we provide evidence to substantiate the claim that *H. lacustris* cysts reproduce via multiple fission. On top of these findings, as schematically shown in Fig. 2F, we conclude that NSC of *H. lacustris* possess the independent and complete life history and propose that *H. lacustris* encompasses two distinct phenotype-dependent life histories: one with the growth-devoting life history (SVC-based) and another specialized for the resistance-dedicated life history (NSC-based). Moreover, the life history of NSC aligns well with the standards of K strategy theory, characterized by a small population size but high stress-resistant offspring. We thus believe that our results provide experimental evidence for the shift between r and K strategies in an isogenic microalga.

### Physiological features of r and K strategists

To gain more physiological details of each strategist, we evaluated and contrasted their corresponding physiological parameters and stress resistance-related traits. We embarked on examining photosynthetic performance of each strategist, one of the most pivotal functions of photosynthetic organisms including microalgae ^39^. Since NSC (K strategists) trade growth for stress resistance, it is reasonable to find that the photosynthesis rate in SVC (r strategists) was significantly higher than that exhibited by K strategists (Fig. 3A, t test, *p* < 0.001). Meanwhile, another growth-related trait, nutrient uptake capability, which is the main nitrogen source for nucleotide and protein biosynthesis and required for cell growth and division ^40^, was examined. Consistent with their photosynthetic performance (Fig. 3A), as shown in Fig. 3B and 3C, r strategists consumed approximately three times as much nitrate as K strategists (*t* test, *p* < 0.01), indicating that r strategists prefer to assimilate a greater quantity of growth-related nutrients, thereby partially elucidating their ability to proliferate and divide at a faster rate compared to K strategists.

**Fig. 3.**
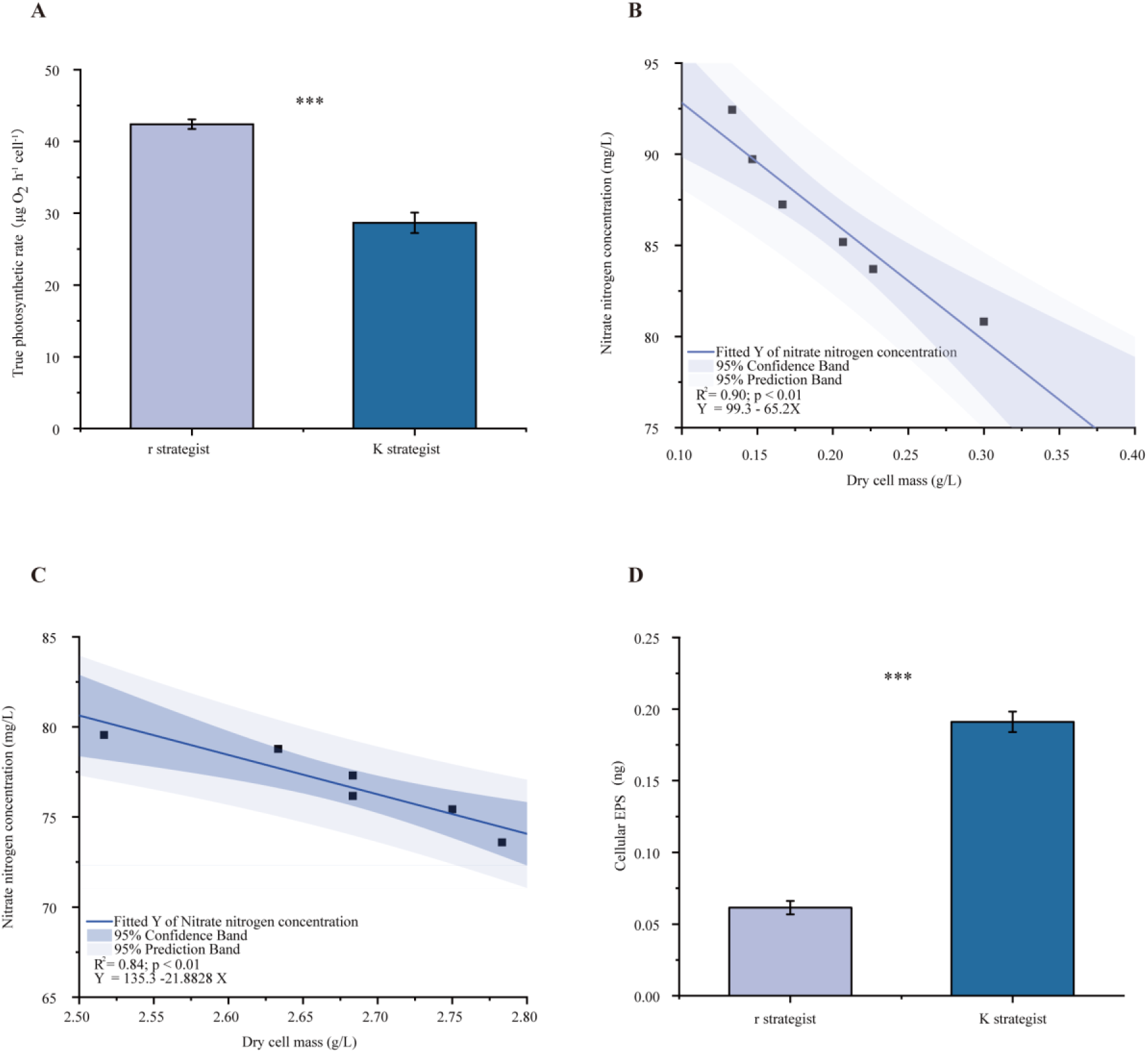
Physiological characteristics of each strategist. (**A**) True photosynthetic rate. (**B**) Nitrate concentration contribution to the dry weight of the SVC (r strategist). (**C**) Nitrate concentration contribution to the dry weight of the NSC (K strategist). (**D**) Cellular EPS content. The error bars indicate the standard deviation of the triplicate values. N = 3 independent biological replicates. Statistical significance was calculated by Welch’s *t* test for pairwise comparisons of two treatments. Significance: * (*p* < 0.05), ** (*p* < 0.01), *** (*p* < 0.001).

In addition, we quantified cellular protein and starch contents. As illustrated in Fig. S4A and S4B, for K strategists, both cellular protein (*t* test, *p* < 0.01) and starch contents (*t* test, *p* < 0.001) were significantly higher than that of r strategists. To mitigate potential confounding effects arising from the increase in dry weight, we analysed the protein/dry weight ratio and the starch/dry weight ratio. The r strategists demonstrated a markedly elevated protein/dry weight ratio, approximately 2.5 times higher (Fig. S4C, *t* test, *p* < 0.001), in contrast, their starch/dry weight ratio was notably reduced, constituting 59.5% of that observed of K strategists (Fig. S4D, *t* test, *p* < 0.001). These comparisons provide valuable insights into the distinct energy utilization tactics adopted by the two strategists, given the pivotal roles of protein in exponential growth rate and starch in stress response^41,42^.

Since the ratio of carbon to nitrogen (C/N) is another important indicator for comprehensively assessing cellular nutrient status, i.e., growth vigour and stress resistant ^43^, we analysed the C/N ratio of each strategist. In general, higher C/N ratios signify high defence capabilities, while lower C/N ratios are indicative of reduced defence potential ^44^. Results illustrate a markedly elevated C/N ratio to 6.5 for K strategists compared with r strategists (Fig. S4E, *t* test, *p* < 0.001), further explaining why K strategists are defence-dedicated from a metabolic perspective.

Finally, exopolysaccharides (EPS) production, another indicator of cellular stress resistance^45^, was evaluated. As shown in Fig. 3D, K strategists produced significantly higher levels (3.1-fold higher) of EPS than that of r strategists (*t* test, *p* < 0.001), which partially explains their higher survival under stressful conditions, such as drought (Fig. 1F).

### Transcriptomic patterns of r and K strategists

RNA-sequencing (RNA-Seq) analysis was employed to understand the underlying molecular background of each strategist and differentially expressed genes (DEGs) of respective cell types were examined. In general, 7330 genes of K strategists showed statistically significant changes in mRNA level compared to that of r strategists (adjusted *p* value < 0.05), with the transcript level of 3750 genes increased and 3580 genes decreased. According to KEGG annotation results, all DEGs were classified into several functional categories, including photosynthesis-antenna protein, nitrogen metabolism, ATP-binding cassette (ABC) transporters, fatty acid elongation, pyruvate metabolism, starch metabolism, ribosome biosynthesis (Fig. S5, top 20 pathways). Since the key finding of this study is that K strategists can reproduce under stress, special attention was paid to genes involved in cell division, e.g., mitosis and ribosome biosynthesis genes.

Intriguingly, in stark contrast to their limited growth and division over a short time, K strategists showed markedly higher reproduction-involved activities at the molecular level compared to the typically dividing r strategists. This observation provides further evidence supporting the phenomenon of multiple fission (Fig. 2B). Specifically, cyclin-dependent kinases genes and cyclin complexes genes, which activate DNA replication, chromosomal segregation, and mitotic steps, were first examined ^46^. Twelve DEGs involved in mitosis were identified in total, among which nine DEGs showed higher expression levels than that of r strategists (Fig. 4A), suggesting an active cell division of K strategists. Simultaneously, we analysed ribosome biosynthesis genes, which serve as another molecular indicator of cell growth and division ^47^. As shown in Fig. 4B, compared to that of K strategists, an upregulated pattern of ribosome biosynthesis DEGs of r strategists were identified, in which 33 out of 41 DEGs (KEGG map03008) showed an upregulated pattern. The expression patterns of genes involved in mitosis and ribosome biosynthesis of K strategists provide extra evidence supporting our observation that K strategists can reproduce, and explaining the molecular mechanisms underlying their reproductive capabilities.

**Fig. 4.**
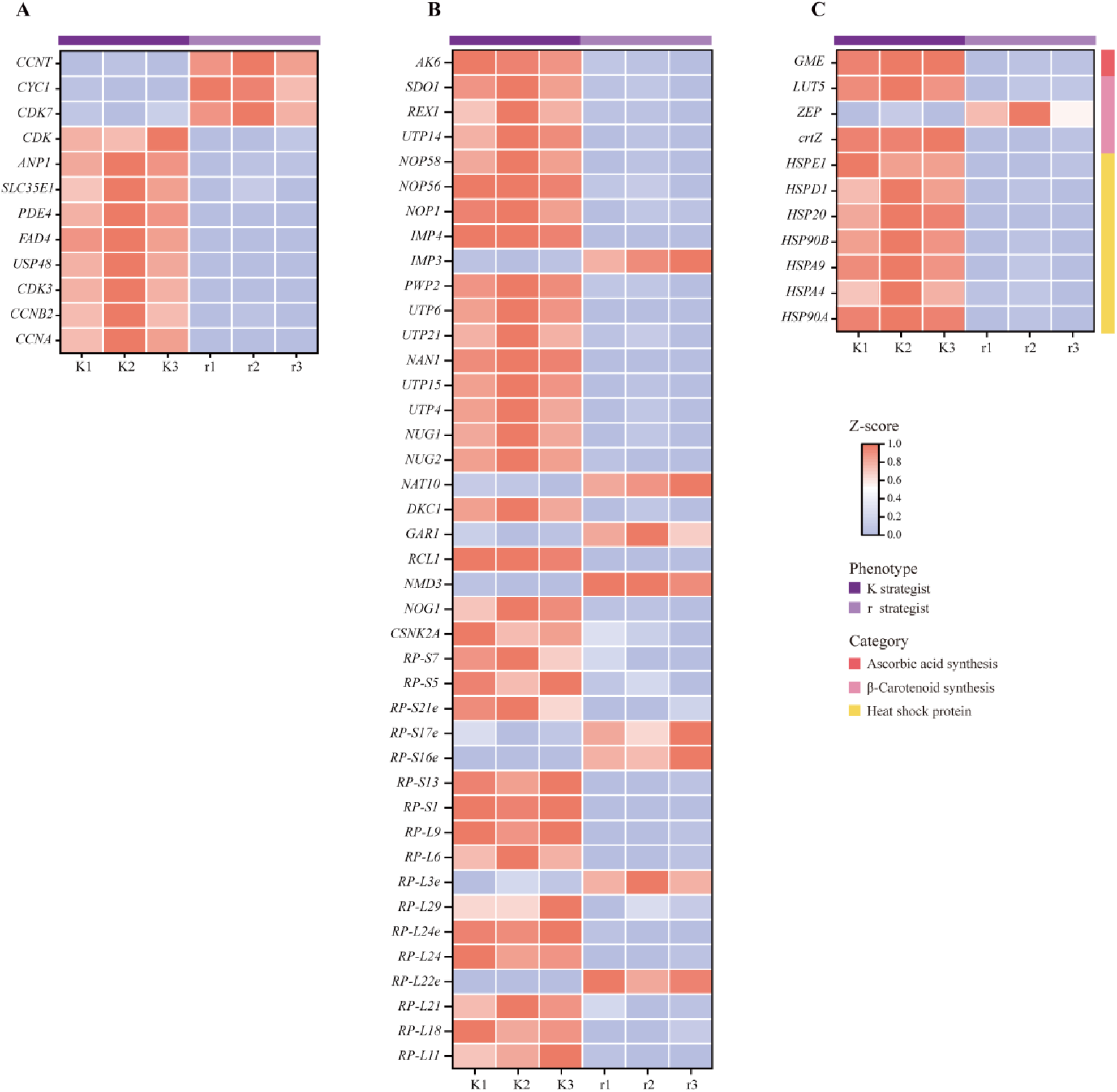
DEGs enrichment analysis of each strategist. (**A**) Heatmap analysis of DEGs involved in mitosis. (**B**) Heatmap analysis of DEGs involved in ribosome biosynthesis. (**C**) Heatmap analysis of DEGs associated with cellular stress resistance, including biosynthesis of ascorbic acid (red), heat shock proteins (yellow) and of β-carotene biosynthesis (pink). The heatmap data is normalised, the blue colour represents low gene expression and the red colour represents high gene expression. The light purple bars represent the r strategist (r), while the dark purple bars represent the K strategist (K). Gene names are shown as KO names. Differentially expressed genes (DEGs) were identified by using DESeq2 with q value ≤ 0.05, accompanied by an absolute value of log2FC (log2 fold change) ≥ 1.

Furthermore, to better understand the underlying mechanisms of the high stress resistance of K strategists, on the one hand, we focused on the genes associated with the biosynthesis of antioxidants, e.g., vitamins ^48^ and heat shock proteins ^49^, which had been shown to play pivotal roles in stress resistance. As illustrated in Fig. 4C, DEGs involved in vitamin C (ascorbic acid) and heat shock proteins in K strategists both showed an upregulated pattern than that of r strategists. On the other hand, genes related to the biosynthesis of β-carotene, which aids in filtering high light irradiance ^50^ and thereby alleviates high light stress, were evaluated. Here, we found most DEGs (10 out of 11 genes) involved in β-carotene biosynthesis showed an upregulated pattern in K strategists than that in r strategists. These transcriptomic patterns collectively elucidate the greater stress resistance observed in K strategists compared to r strategists (Fig. 1E, Fig. 1F).

### Traits of substance and energy metabolism of r and K strategists

Substance and energy metabolism are among the most crucial processes for all living organisms ^51^, which are also keystones for cellular growth and defence. Accordingly, the transcriptomic discrepancies in substance and energy metabolism between each strategist were contrasted ^52^. Of note, to show a more complete gene expression pattern in a particular pathway, all annotated genes (including none-DEGs) were included in the detailed analyses.

In terms of substance metabolism, r strategists exhibited a superior nitrogen assimilation capacity compared to K strategists. This is evidenced by the upregulation of all genes associated with nitrogen metabolism (KEGG pathway map00910) in r strategists relative to K strategists (Fig. 5). These findings align with the results obtained from the nitrogen consumption and protein/dry weight ratio assays (Fig. 3B & Fig. S4C). In contrast, as s illustrated in Fig. 5, the expression of carbon assimilation genes (starch and sucrose metabolism, KEGG map00500) were generally upregulated in K strategists (five out of six annotated genes), compared to that of r strategists. This pattern aligns with higher cellular starch content and elevated C/N ratio in K strategists (Fig. S4D and Fig. S4E).

**Fig. 5.**
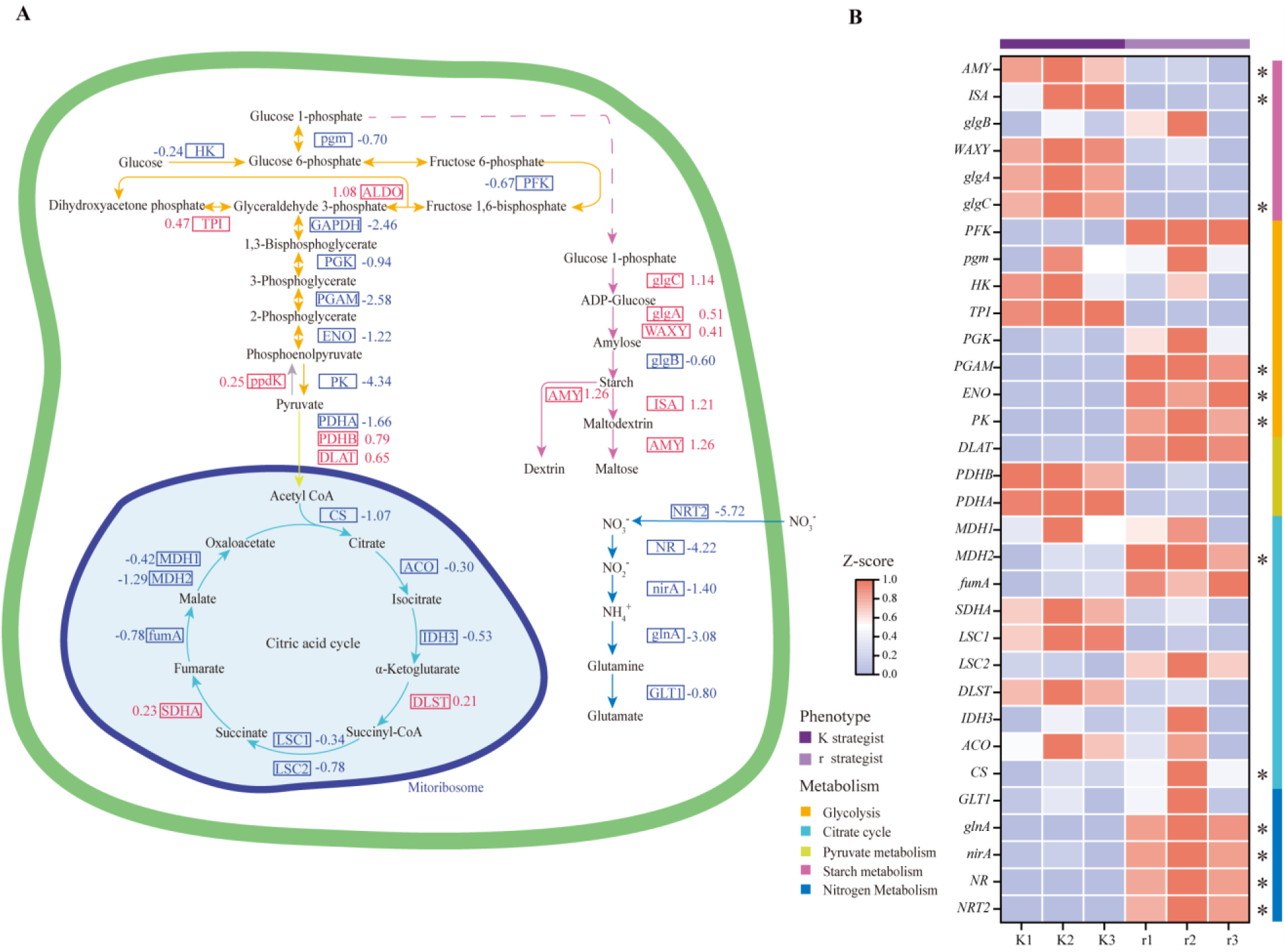
Different substance and energy metabolic traits of each strategist. (**A**) Schematic presentation of energy metabolism-related pathways, including glycolysis (orange), citrate cycle (light indigo), pyruvate metabolism (yellow), starch metabolism (pink), and nitrogen assimilation (dark indigo). Compared with that of the r strategist (r), upregulated (red) and downregulated (blue) genes of the K strategist (K) are illustrated. The dotted lines adorned with arrows serve to illustrate the subsequent trajectory of the substance flow. (**B**) Heatmap analysis of genes involved in substance and energy metabolism. The heatmap data is normalised, the blue colour represents low gene expression and the red colour represents high gene expression. The different colours of the bar on the right show the pathways, including glycolysis (orange), citrate cycle (light indigo), pyruvate metabolism (yellow), starch metabolism (pink), and nitrogen assimilation (dark indigo). The light purple bar represents the r strategist, while the dark purple bar represents the K strategist. Gene names are shown as KO names. Differentially expressed genes (DEGs) are marked with asterisks, which were identified by using DESeq2 with q value ≤ 0.05, accompanied by an absolute value of log2FC (log2 fold change) ≥ 1.

To understand the energetic traits of r and K strategists, we evaluated transcriptomic patterns of genes involved in photosynthesis and respiration, two key energy metabolisms in photoautotrophic microalgae ^53–55^. More specifically, genes related to photosynthesis-antenna protein, photosynthesis, glycolysis and tricarboxylic acid cycle (TCA) were evaluated. In general, r strategists exhibit a superior capacity for energy capture and production compared to K strategists, as indicated by their upregulated gene expression levels in key energy-related pathways. In detail, r strategists showed upregulated gene expression levels in: 1) the photosynthesis - antenna protein (9 out of 9, map00196, Fig. S6) and photosynthesis pathway (12 out of 14, map00195, Fig. S6); 2) glycolysis (8 out of 9 annotated genes, map00010, Fig. 5) and TCA cycle (8 out of 10 genes, map00020, Fig. 5).

Collectively, for substance metabolism, K strategists prefer carbon assimilation, leading to the storage of energetic compounds (starch). This accumulation is essential for enhancing stress endurance. In contrast, r strategists tend to assimilate higher levels of nitrogen, which facilitates the synthesis of protein and nucleotides, building blocks for cell growth and division. For energy metabolism, r strategists exhibit greater efficiency in energy capture and generation compared to K strategists, implying their preference for energy-consuming growth and reproduction.

## Discussion

For *H. lacustris*, it has been well-known that SVC can transform to NSC along with the deterioration of external environments, however, to our knowledge, NSC are almost universally assumed to be a dormant state that are incapable of reproduction under stress ^18,19,25,33^. In this study, by integrating evidence from cellular size comparison, direct microscopic observation as well as the transcriptomic patterns of reproduction-involved genes, we show that NSC can reproduce in the mode of multiple fission, dividing into eight daughter cells. Notably, the observed multiple fission occurred in old cultures (adverse conditions), and the corresponding daughter cysts remained non-mobile and stress resistant, which is in stark contrast to cyst germination ^56^, herein daughter cells regain mobility, normal growth and reproduction (binary fission) upon transferring into favourable environments, such as fresh medium.

Since NSC can reproduce under adverse conditions, it thus can be concluded that they own a complete life history. Moreover, given their low growth rate (barely detectable over a short period, Fig. 1D) and stress-resistant daughter cells at birth (Fig. 2E), it is reasonable to define NSC as K strategists. Furthermore, similar to many other microbes, SVC of *H. lacustris*, which showed rapid growth and division (Fig. 1D) but were sensitive to external stresses (Fig. 1E & Fig. 1F), generally meet the criteria of r strategists, we therefore present experimental evidence to conclude the shift between r and K strategies in the unicellular microalga *H. lacustris*.

Different from other organisms that either engage in r strategy or K strategy, the employ of both strategies can confer *H. lacustris* fitness advantages under both favourable and adverse conditions. When external environments are favourable, *H. lacustris* carries out the r strategy, favouring reproduction and leading to a high quantity of swimming daughter cells (still r strategists). When environments turn unfavourable, *H. lacustris* activates the K strategy program, reproducing a limited number of stress-resistant offspring.

In nature, such a life history strategy shift (from r to K) guarantees the prosper and survival of *H. lacustris* populations, especially under fluctuating natural environments. Indeed, different from many other microalgal species living in more stable open waters, the natural habitats of *H. lacustris* are often small and temporary aquatic habitats ^20,21^, tending to change quickly and dramatically, posing challenges to dwelling organisms. Under such conditions, for *H. lacustris*, the engagement in both r and K strategies could serve as an essential and effective adaptation strategy, it confers *H. lacustris* growth and reproduction advantages to maintain growth under both unfavourable and favourable conditions over other microalgal species, that are sensitive to the challenge of environmental distributions. Moreover, since cyst formation is a common stress-response strategy identified in many microalgal species ^17^, it is interesting to explore whether these microalgal cysts can also reproduce. In the context of our results, the reproduction capability of microalgal cysts under stresses might be overlooked.

In summary, in contrast to the conventional idea that microalgal cysts cannot reproduce under stress (thus distinct from cyst germination when environments turn favourable), here we provide evidence showing that *H. lacustris* cysts (NSC) can reproduce in the mode of multiple fission under continuous stress. In this case, similar to SVC that possess a complete life history, these NSC are concluded to have a complete life history as well. Since *H. lacustris* cysts reproduce less but higher stress-resistant offspring than vegetative cells, we thus define their life history as K strategy. Therefore, based on the fact that growth-dedicated SVC and stress-resistant NSC of *H. lacustris* employ the r strategy and K strategy, respectively, we propose the shift between r and K strategies in this isogenic strain, offering new perspectives to understand microalgal survival and adaptation strategies in a challenging world.

## Materials and Methods

### Strain and culture conditions

*Haematococcus lacustris* FACHB-712 (the core strain of this study), FACHB-797, FACHB-871 and FACHB-1928 were obtained from the Freshwater Algae Culture Collection at the Institute of Hydrobiology, Wuhan, China. Stock culture was maintained in 250 mL BBM ^57^ in Erlenmeyer flasks at 23±1 °C under 12h/12 h light/dark cycles with a light intensity of 18 μmol photons m^−2^ s^−1^ unless otherwise specified, all experiments were conducted under the same conditions (standard conditions).

### Growth experiment and phenotypic transformation

Triplicate *H. lacustris* populations starting with 100% of SVC with an initial cell density of approximately 2.4 × 10^5^ cells/mL were grown under standard conditions without external stresses over a 23-day period. Cell density and cellular size of different phenotypes, based on their swimming capability ^22^, were recorded every other day. For cell number and cellular size quantification, images of 100 µL culture, which was first fixed with 5 μL Lugol’s solution, were captured under an inverted microscope (Primo Vert, Carl Zeiss, Germany) with a plankton counting chamber. Images were then processed with the image recognition algorithm (Mask R-CNN, a computer vision algorithm performing instance segmentation ^58,59^) to obtain cell density and cellular size data. For each replicate at each time point, at least 1000 cells were measured.

### Separation of SVC and NSC

When grown in BBM under standard conditions, SVC will gradually lose mobility transforming to NSC and settle to the bottom of the culture flask after two months and thus can be collected for further experiments. In contrast, SVC can be obtained by transferring cells (either swimming, non-swimming or mixed phenotype) into fresh medium with a low cell density, here all cells can remain mobile or regain mobility in a few days.

### Growth under standard or stressful conditions

To explore the advantages of each phenotype, the growth performance of the swimming and non-swimming phenotypes cultured under standard or stressful conditions was examined. For experiments under standard conditions, SVC were inoculated in fresh BBM medium (with a starting density of approximately 1.3 × 10^5^ cells/mL), whereas the supernatants of NSC (2-month-old) were first harvested and NSC were inoculated in the old culture supernatants maintaining consistency with the concentration of the original culture system (with a starting density of approximately 3.5×10^5^ cells/mL), avoiding the transition from NSC to SVC. Cell number and cellular dry weight was quantified daily, and the experiment lasted for 6 days, conducted in triplicate.

High light irradiation and drought were used as external stresses to evaluate the stress resistance of each phenotype. For the high light irradiation exposure, triplicate populations of SVC and NSC (starting with 3.6×10^5^ cells/mL) were grown under a light intensity of 145 μmol photons m^−2^ s^−1^ for 3 days, and cell densities were recorded on the first day and the last day of the experiment. For drought stress, BBM agar plates (2% agar) were prepared to create an artificial drought environment, and cells were evenly spread using the spread plate method. Based on the number of colonies, survivors of each treatment were enumerated after 3 weeks. All experiments were conducted in triplicate.

### Growth performance of daughter cells of each phenotype

To explore the benefits of each life history strategy, the growth performance of daughter cells from each strategy under the stressful condition was examined. To this end, we first collected the daughter cells of r strategists and K strategist, respectively. For SVC, since populations are active in reproduction, cells of the exponential growth phase (day 3) are considered as daughter SVC. For daughter cells of NSC, since cells generally increase in size during the transition from the swimming to non-swimming phenotype, NSC smaller than the average size of SVC is defined as daughter NSC. We therefore collected smaller NSC using a mesh filter to remove bigger cells. To conduct a more comprehensive exploration of stress resistance, salinity (100 mM, 200 mM, 300 mM NaCl) was included as the external stress, and the performance of collected daughter cells (starting with 5 × 10^4^ cells/mL) grown in the presence of NaCl was recorded. This experiment was carried out in triplicate and ran for 7 days, cell numbers were checked on day 1 and day 7, and survival rates were calculated.

### Cellular size dynamics of different *H. lacustris* strains

Three other *H. lacustri*s strains (FACHB-797, Hubei province, China; FACHB-871, Jiangxi province, China; FACHB-1928, Yunnan province, China) were used for the experiment. SVC and NSC were separated and cellular size of each phenotype was then quantified as described above.

### Microscopy

Microscopic images were taken using an inverted fluorescence microscope (Primo Vert, Carl Zeiss, Germany). 1mL SVC and NSC were centrifuged at 800 × g for 10 min, respectively. Supernatants were discarded, and cells were washed gently with 0.1 M phosphate buffer solution (PBS, pH 7.4) three times. Then 100 µL culture was fixed with 5 μL Lugol’s solution, which later was used for observation and images capture.

### Nitrate quantification

Residual nitrate content of the culture medium of two phenotypes was measured daily. Supernatants of triplicate samples of each time point were collected, and residual nitrate was quantified using the Auto Discrete Analyzer (Cleverchem Anna, Dechem-Tech, Germany). In particular, nitrate nitrogen concentrations were measured using the cadmium-zinc reduction method ^60^.

### Cellular total carbon and total nitrogen quantification

Cells of each phenotype were centrifuged and freeze-dried, then 5 mg sample was wrapped in the tinfoil. Collected samples were oxidised in the reaction tube to produce N2 and CO_2_ using MACRO cube elemental analyser (Elmentar, Germany), and then the TC and TN were measured, as described by Yun Yan ^61^. C% and N% were calculated by detecting the peak area and comparing it with a standard sample, i.e., sulphate.

### Quantification of cellular dry weight, protein, starch and EPS

For the determination of cellular dry weight, 5 mL of respective samples were collected and centrifuged, then, cell pellets were washed three times with 0.1 M phosphate buffer solution (PBS, pH 7.4) using a pre-dried PES sterile syringe filter (25 mm in diameter, 0.45 μm of pore size, Tianjin Jinteng). Collected cell pellets were then dried for six hours at 50 °C in an oven (Yiheng, Shanghai, China) and cooled for three hours to room temperature before weighing.

For cellular protein quantification, 1 mL sample of each treatment was harvested by centrifugation at 800 × g for 10 min. Collected cells were then hydrolysed with 1 M NaOH in a water bath at 70 °C for 15 min, and protein content was determined with a spectrophotometer (Infinite® M200 PRO, TECAN, America) at 562 nm using an enhanced BCA protein assay kit (ZT10012, ZETA, America), as previously described ^62^.

Cellular starch was quantified according to a published method with some modifications ^63^. 2 mL of sample was ground with an automatic sample fast grinder (Shanghai Jingxin, China), 1 mL of which was used for further quantification. After centrifugation at 800 × g for 10 min, the pigments in the pellets were first extracted with 4 mL of 80% ethanol for 15 min at 70 °C, the process was repeated until the pellets were colourless. For complete hydrolysis of the starch, 1 mL of 52% perchloric acid was added to the precipitate, stirred for 45 min at 25 °C and centrifuged. Then 0.2 mL of the extract was cooled to 0 °C; 1 mL of anthrone solution [0.1 g of anthrone in 50 mL of 82% (v/v) H2SO4] was added and stirred. The mixture was kept in a water bath at 100 °C for 8 min and then cooled to 0 °C immediately. After that the samples were kept at room temperature for an additional 10 min. The absorbance was measured at 625 nm. Calibration was performed simultaneously with glucose as the standard.

The exopolysaccharides (EPS) were extracted using a heating method ^64^. Briefly, 2 mL of algal cells were collected and centrifuged at 800 × g for 10 min. The remaining algal pellet was washed with 0.1 M phosphate buffer solution (PBS, pH 7.4) three times, then resuspended in an equal volume of 0.05% NaCl solution. This suspension was heated in a water bath at 60 °C for 15 min and subsequently centrifuged at 6500 × g for 20 min at 4 °C. The supernatant was filtered through a 0.22 μm membrane. The EPS content was determined using a total organic carbon analyser (TOC-L, Shimadzu, Japan).

### True photosynthesis rate estimation

YZQ-201A photosynthetic instrument (Yizongqi Technology Co., Beijing, China) was used to test the true photosynthesis rate with luminescent DO sensors (LDO). The real-time suspension oxygen concentration of the sample was measured based on the quenching of luminescence in the presence of oxygen. The conditions are as follows: 180 s stabilisation time, 15 mL sample volume, 18 μmol m^-2^ s^-1^ light intensity. The oxygen consumption rate was measured under dark for 300 s at 20 °C and the oxygen production rate was measured for 300 s in light (RGB light source). The true photosynthesis rate was calculated in the following formulas:

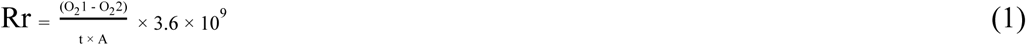

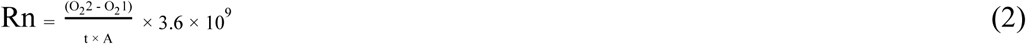

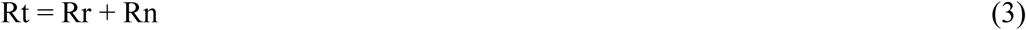

Where O_2_1 is the initial oxygen concentration (mg/L), O_2_2 is the end oxygen concentration (mg/L), t is the detection time (s), A is the cell number (mL^-1^), Rr is the respiration rate (μg O_2_ h^-1^ cell^-1^) measured in dark period, Rn is the net photosynthetic rates (μg O_2_ h^-1^ cell^-1^) measured in light period, Rt is the true photosynthetic rates (μg O_2_ h^-1^ cell^-1^) ^65^.

### Transcriptomics

For sample preparation, 50 mL of each cell types was centrifuged for 15 min at 800 × g at 4 °C using RNase-free tubes and the supernatant was removed. 50 mL of PBS (0.1 M, pH 7.4) was added, and cell pellets were washed three times. Then, the cell pellets were frozen in liquid nitrogen for at least 30 min. In total, both cell types were prepared in triplicate. Next, RNA samples were extracted and sent to Shanghai Majorbio Bio-pharm Technology Co., Ltd. (Shanghai, China) to construct an RNAseq transcriptome library.

For data analysis, the Majorbio Cloud online platform (www.majorbio.com) was used. After raw reads were quality-trimmed and filtered (SeqPrep and Sickle), clean reads were aligned against the *H. lacustris* genome (https://www.ncbi.nlm.nih.gov/datasets/genome/GCA_030144725.1/) with HISAT2. Mapped reads were assembled by using StringTie ^66^. Transcript expression levels were calculated by using the transcripts per million reads method (TPM). Differentially expressed genes (DEGs) were identified by using DESeq2 with q value ≤ 0.05, accompanied by an absolute value of log2FC (log2 fold change) ≥ 1. Subsequently, Kyoto Encyclopedia of Genes and Genomes (KEGG) enrichment was analysed by Goatools and KOBAS.

### Statistics

All statistical analyses were performed in OriginPro. Statistical significance was calculated by Welch’s *t* test for pairwise comparisons of two treatments (*p* value < 0.05). Significance: * (*p* < 0.05), ** (*p* < 0.01), *** (*p* < 0.001).

## Acknowledgements

We thank Qi Yan for the help with technical support and discussions and Kebi Wu for assistance with algal cultures and Wenjie Zhang for logistical and sampling support. We also thank W. Wang for NMR data collection.

## Finding

This work was supported by National Natural Science Foundation of China (41976126)

The S&T Projects of Shenzhen Science and Technology Innovation Committee (KCXFZ20230731093402005, RCJC20200714114433069, and SGDX20220530111204028)

## Author contributions

Conceptualization: LL, ZHC, ST

Methodology: LL, ZHC, ST

Investigation: LL, YQL, YLZ

Visualization: LL, BYZ, DL

Supervision: ZHC

Writing—original draft: LL, ST

Writing—review & editing: LL, ST, LYQ, BYZ, YLZ, DL, JZ, ZHC

## Data and materials availability

All data are available in the main text or the supplementary materials. The raw data of RNA-Seq are available from NCBI’s SRA database (http://www.ncbi.nlm.nih.gov) under BioProject PRJNA1157585. KEGG database (https://www.kegg.jp/) was used for functional enrichment analyses. Source data are provided with this paper.

## References

1. He, Z., Webster, S., and He, S.Y. (2022). Growth–defense trade-offs in plants. Current Biology 32, R634–R639. 10.1016/j.cub.2022.04.070.

2. Smith, W.P.J., Wucher, B.R., Nadell, C.D., and Foster, K.R. (2023). Bacterial defences: mechanisms, evolution and antimicrobial resistance. Nat Rev Microbiol 21, 519–534. 10.1038/s41579-023-00877-3.

3. Lean, J.L., and Rind, D.H. (2008). How natural and anthropogenic influences alter global and regional surface temperatures: 1889 to 2006. Geophysical Research Letters 35. 10.1029/2008GL034864.

4. Omer, A., Zhuguo, M., Zheng, Z., and Saleem, F. (2020). Natural and anthropogenic influences on the recent droughts in Yellow River Basin, China. Science of The Total Environment 704, 135428. 10.1016/j.scitotenv.2019.135428.

5. Andrews, J.H., and Harris, R.F. (1986). r- and K-Selection and Microbial Ecology. In Advances in Microbial Ecology Advances in Microbial Ecology., K. C. Marshall, ed. (Springer US), pp. 99–147. 10.1007/978-1-4757-0611-6_3.

6. Hengeveld, R. (2002). The Theory of Island Biogeography. Acta Biotheoretica 50, 133–136. 10.1023/A:1016393430551.

7. Stearns, S.C. (1976). Life-History Tactics: A Review of the Ideas. The Quarterly Review of Biology 51, 3–47. 10.1086/409052.

8. Malik, A.A., Martiny, J.B.H., Brodie, E.L., Martiny, A.C., Treseder, K.K., and Allison, S.D. (2020). Defining trait-based microbial strategies with consequences for soil carbon cycling under climate change. ISME J 14, 1–9. 10.1038/s41396-019-0510-0.

9. Richardson, B.J. (1975). r and K selection in kangaroos. Nature 255, 323–324. 10.1038/255323a0.

10. Schmidt, W., and Briibach, M. (1993). Plant distribution patterns during early succession on an artificial protosoil. Journal of Vegetation Science 4, 247–254. 10.2307/3236111.

11. Yin, Q., Sun, Y., Li, B., Feng, Z., and Wu, G. (2022). The r/K selection theory and its application in biological wastewater treatment processes. Science of The Total Environment 824, 153836. 10.1016/j.scitotenv.2022.153836.

12. Sinervo, B., Svensson, E., and Comendant, T. (2000). Density cycles and an offspring quantity and quality game driven by natural selection. Nature 406, 985–988. 10.1038/35023149.

13. Cheng, Y., Hubbard, C.G., Zheng, L., Arora, B., Li, L., Karaoz, U., Ajo-Franklin, J., and Bouskill, N.J. (2018). Next generation modeling of microbial souring – Parameterization through genomic information. International Biodeterioration & Biodegradation 126, 189–203. 10.1016/j.ibiod.2017.06.014.

14. Pianka, E.R. (1970). On r- and K-Selection. The American Naturalist 104, 592–597. 10.1086/282697.

15. Fierer, N., Bradford, M.A., and Jackson, R.B. (2007). Toward an ecological classification of soil bacteria. Ecology 88, 1354–1364. 10.1890/05-1839.

16. Fierer, N., Lauber, C.L., Ramirez, K.S., Zaneveld, J., Bradford, M.A., and Knight, R. (2012). Comparative metagenomic, phylogenetic and physiological analyses of soil microbial communities across nitrogen gradients. ISME J 6, 1007–1017. 10.1038/ismej.2011.159.

17. Jones, S.E., and Lennon, J.T. (2010). Dormancy contributes to the maintenance of microbial diversity. Proceedings of the National Academy of Sciences of the United States of America 107, 5881–5886. 10.1073/pnas.0912765107.

18. Figueroa, R.I., Estrada, M., and Garcés, E. (2018). Life histories of microalgal species causing harmful blooms: Haploids, diploids and the relevance of benthic stages. Harmful Algae 73, 44–57. 10.1016/j.hal.2018.01.006.

19. Lennon, J.T., and Jones, S.E. (2011). Microbial seed banks: the ecological and evolutionary implications of dormancy. Nat Rev Microbiol 9, 119–130. 10.1038/nrmicrO2504.

20. Chekanov, K., Fedorenko, T., Kublanovskaya, A., Litvinov, D., and Lobakova, E. (2019). Diversity of carotenogenic microalgae in the White Sea polar region. FEMS Microbiology Ecology, fiz183. 10.1093/femsec/fiz183.

21. Kublanovskaya, A., Baulina, O., Chekanov, K., and Lobakova, E. (2020). The microalga Haematococcus lacustris (Chlorophyceae) forms natural biofilms in supralittoral White Sea coastal rock ponds. Planta 252, 37. 10.1007/s00425-020-03438-7.

22. Tang, S., Liu, Y., Zhu, J., Cheng, X., Liu, L., Hammerschmidt, K., Zhou, J., and Cai, Z. (2024). Bet hedging in a unicellular microalga. Nat Commun 15, 2063. 10.1038/s41467-024-46297-6.

23. Ginzberg, M.B., Kafri, R., and Kirschner, M. On being the right (cell) size. CELL BIOLOGY.

24. Kobayashi, M., Kurimura, Y., Kakizono, T., Nishio, N., and Tsuji, Y. (1997). Morphological changes in the life cycle of the green alga Haematococcus pluvialis. Journal of Fermentation and Bioengineering 84, 94–97. 10.1016/S0922-338X(97)82794-8.

25. Wang, N., Guan, B., Kong, Q., and Duan, L. (2018). A semi-continuous cultivation method for Haematococcus pluvialis from non-motile cells to motile cells. J Appl Phycol 30, 773–781. 10.1007/s10811-017-1337-6.

26. Rengefors, K., Karlsson, I., and Hansson, L. (1998). Algal cyst dormancy: a temporal escape from herbivory. Proc. R. Soc. Lond. B 265, 1353–1358. 10.1098/rspb.1998.0441.

27. Richardson, B., Orcutt, D.M., Schwertner, H.A., Martinez, C.L., and Wickline, H.E. (1969). Effects of Nitrogen Limitation on the Growth and Composition of Unicellular Algae in Continuous Culture. Appl Microbiol 18, 245–250.

28. Giraldo, N.D., Correa, S.M., Arbeláez, A., Figueroa, F.L., Ríos-Estepa, R., and Atehortúa, L. (2021). Reducing self-shading effects in Botryococcus braunii cultures: effect of Mg2+ deficiency on optical and biochemical properties, photosynthesis and lipidomic profile. Bioresour. Bioprocess. 8, 33. 10.1186/s40643-021-00389-z.

29. Hadj-Romdhane, F., Zheng, X., Jaouen, P., Pruvost, J., Grizeau, D., Croué, J.P., and Bourseau, P. (2013). The culture of Chlorella vulgaris in a recycled supernatant: Effects on biomass production and medium quality. Bioresource Technology 132, 285–292. 10.1016/j.biortech.2013.01.025.

30. Tang, Y.Z., Gu, H., Wang, Z., Liu, D., Wang, Y., Lu, D., Hu, Z., Deng, Y., Shang, L., and Qi, Y. (2021). Exploration of resting cysts (stages) and their relevance for possibly HABs-causing species in China. Harmful Algae 107, 102050. 10.1016/j.hal.2021.102050.

31. Ellegaard, M., and Ribeiro, S. (2018). The long-term persistence of phytoplankton resting stages in aquatic ‘seed banks.’ Biological Reviews 93, 166–183. 10.1111/brv.12338.

32. Dahms, H.-U. (1995). Dormancy in the Copepoda ? an overview. Hydrobiologia 306, 199– 211. 10.1007/BF00017691.

33. Boussiba, S. (2000). Carotenogenesis in the green alga Haematococcus pluvialis: Cellular physiology and stress response. Physiol Plant 108, 111–117. 10.1034/j.1399-3054.2000.108002111.x.

34. Zachleder, V., Bišová, K., and Vítová, M. (2016). The Cell Cycle of Microalgae. In The Physiology of Microalgae, M. A. Borowitzka, J. Beardall, and J. A. Raven, eds. (Springer International Publishing), pp. 3–46. 10.1007/978-3-319-24945-2_1.

35. Ivanov, I.N., Vítová, M., and Bišová, K. (2019). Growth and the cell cycle in green algae dividing by multiple fission. Folia Microbiol 64, 663–672. 10.1007/s12223-019-00741-z.

36. Wayama, M., Ota, S., Matsuura, H., Nango, N., Hirata, A., and Kawano, S. (2013). Three-Dimensional Ultrastructural Study of Oil and Astaxanthin Accumulation during Encystment in the Green Alga Haematococcus pluvialis. PLoS ONE 8, e53618. 10.1371/journal.pone.0053618.

37. Lakshmi Narasimhan, A., Lee, N., Kim, S., Kim, Y.-E., Christabel, C., Yu, H., Kim, E.-J., and Oh, Y.-K. (2024). Enhanced astaxanthin production in Haematococcus lacustris by electrochemical stimulation of cyst germination. Bioresource Technology 411, 131301. 10.1016/j.biortech.2024.131301.

38. Genovesi, B., Laabir, M., Masseret, E., Collos, Y., Vaquer, A., and Grzebyk, D. (2009). Dormancy and germination features in resting cysts of Alexandrium tamarense species complex (Dinophyceae) can facilitate bloom formation in a shallow lagoon (Thau, southern France). 31.

39. Shimakawa, G., Matsuda, Y., Nakajima, K., Tamoi, M., Shigeoka, S., and Miyake, C. (2017). Diverse strategies of O_2_ usage for preventing photo-oxidative damage under CO_2_ limitation during algal photosynthesis. Sci Rep 7, 41022. 10.1038/srep41022.

40. Stein, L.Y., and Klotz, M.G. (2016). The nitrogen cycle. Current Biology 26, R94–R98. 10.1016/j.cub.2015.12.021.

41. Robyt, J.F. (2008). Starch: Structure, Properties, Chemistry, and Enzymology. In Glycoscience: Chemistry and Chemical Biology, B. O. Fraser-Reid, K. Tatsuta, and J. Thiem, eds. (Springer), pp. 1437–1472. 10.1007/978-3-540-30429-6_35.

42. Utting, S.D. (1985). Influence of nitrogen availability on the biochemical composition of three unicellular marine algae of commercial importance. Aquacultural Engineering 4, 175–190. 10.1016/0144-8609(85)90012-3.

43. Xu, X., Yang, G., Yang, X., Li, Z., Feng, H., Xu, B., and Zhao, X. (2018). Monitoring ratio of carbon to nitrogen (C/N) in wheat and barley leaves by using spectral slope features with branch-and-bound algorithm. Sci Rep 8, 10034. 10.1038/s41598-018-28351-8.

44. Royer, M., Larbat, R., Le Bot, J., Adamowicz, S., and Robin, C. (2013). Is the C:N ratio a reliable indicator of C allocation to primary and defence-related metabolisms in tomato? Phytochemistry 88, 25–33. 10.1016/j.phytochem.2012.12.003.

45. Houston, K., Tucker, M.R., Chowdhury, J., Shirley, N., and Little, A. (2016). The Plant Cell Wall: A Complex and Dynamic Structure As Revealed by the Responses of Genes under Stress Conditions. Frontiers in Plant Science 7. 10.3389/fpls.2016.00984.

46. Kominami, S., Mizuta, H., and Uji, T. (2022). Transcriptome Profiling in the Marine Red Alga Neopyropia yezoensis Under Light/Dark Cycle. Mar Biotechnol 24, 393–407. 10.1007/s10126-022-10121-3.

47. Dai, X., and Zhu, M. (2020). Coupling of Ribosome Synthesis and Translational Capacity with Cell Growth. Trends in Biochemical Sciences 45, 681–692. 10.1016/j.tibs.2020.04.010.

48. Jiménez-Árias, D., García-Machado, F.J., Morales-Sierra, S., Garrido-Orduña, C., Borges, A.A., González, F.V., and Luis Jorge, J.C. (2018). Chapter 7 - Vitamins and Environmental Stresses in Plants. In Plant Metabolites and Regulation Under Environmental Stress, P. Ahmad, M. A. Ahanger, V. P. Singh, D. K. Tripathi, P. Alam, and M. N. Alyemeni, eds. (Academic Press), pp. 145–152. 10.1016/B978-0-12-812689-9.00007-8.

49. Lang, B.J. (2021). The functions and regulation of heat shock proteins; key orchestrators of proteostasis and the heat shock response. Archives of Toxicology. 10.1007/s00204-021-03070-8.

50. Bar, E., Rise, M., Vishkautsan, M., and Arad, S. (Malis) (1995). Pigment and Structural Changes in Chlorella zofingiensis upon Light and Nitrogen Stress. Journal of Plant Physiology 146, 527–534. 10.1016/S0176-1617(11)82019-5.

51. Liu, L., Zeng, X., Zheng, J., Zou, Y., Qiu, S., and Dai, Y. (2022). AHL-mediated quorum sensing to regulate bacterial substance and energy metabolism: A review AHL 介导的群体感应调节细菌物质和能量代谢,综述. Microbiological Research 262, 127102. 10.1016/j.micres.2022.127102.

52. Zhang, C.-C., Zhou, C.-Z., Burnap, R.L., and Peng, L. (2018). Carbon/Nitrogen Metabolic Balance: Lessons from Cyanobacteria. Trends in Plant Science 23, 1116–1130. 10.1016/j.tplants.2018.09.008.

53. Araújo, W.L., Nunes-Nesi, A., Nikoloski, Z., Sweetlove, L.J., and Fernie, A.R. (2012). Metabolic control and regulation of the tricarboxylic acid cycle in photosynthetic and heterotrophic plant tissues. Plant Cell & Environment 35, 1–21. 10.1111/j.1365-3040.2011.02332.x.

54. van Dongen, J.T., Gupta, K.J., Ramírez-Aguilar, S.J., Araújo, W.L., Nunes-Nesi, A., and Fernie, A.R. (2011). Regulation of respiration in plants: A role for alternative metabolic pathways. Journal of Plant Physiology 168, 1434–1443. 10.1016/j.jplph.2010.11.004.

55. Yamori, W. (2016). Chapter 9 - Photosynthesis and respiration. In Plant Factory, T. Kozai, G. Niu, and M. Takagaki, eds. (Academic Press), pp. 141–150. 10.1016/B978-0-12-801775-3.00009-3.

56. Daroch, M. (2016). Astaxanthin-Producing Green Microalga Haematococcus pluvialis: From Single Cell to High Value Commercial Products. Frontiers in Plant Science 7, 28.

57. Marinho, Y.F., Malafaia, C.B., de Araújo, K.S., da Silva, T.D., dos Santos, A.P.F., de Moraes, L.B., and Gálvez, A.O. (2021). Evaluation of the influence of different culture media on growth, life cycle, biochemical composition, and astaxanthin production in Haematococcus pluvialis. Aquacult Int 29, 757–778. 10.1007/s10499-021-00655-z.

58. Bi, H., Song, J., Zhao, J., Liu, H., Cheng, X., Wang, L., Cai, Z., Benfield, M.C., Otto, S., Goberville, E., et al. (2022). Temporal characteristics of plankton indicators in coastal waters: High-frequency data from PlanktonScope. Journal of Sea Research 189, 102283. 10.1016/j.seares.2022.102283.

59. He, K., Gkioxari, G., Dollar, P., and Girshick, R. (2020). Mask R-CNN. IEEE Trans. Pattern Anal. Mach. Intell. 42, 386–397. 10.1109/TPAMI.2018.2844175.

60. Yan, Q., Cheng, T., Song, J., Zhou, J., Hung, C.-C., and Cai, Z. (2021). Internal nutrient loading is a potential source of eutrophication in Shenzhen Bay, China. Ecological Indicators 127, 107736. 10.1016/j.ecolind.2021.107736.

61. Yan, Y., Tian, J., Fan, M., Zhang, F., Li, X., Christie, P., Chen, H., Lee, J., Kuzyakov, Y., and Six, J. (2012). Soil organic carbon and total nitrogen in intensively managed arable soils. Agriculture, Ecosystems & Environment 150, 102–110. 10.1016/j.agee.2012.01.024.

62. Lu, Z., Zheng, L., Liu, J., Dai, J., and Song, L. (2019). A novel fed-batch strategy to boost cyst cells production based on the understanding of intracellular carbon and nitrogen metabolism in Haematococcus pluvialis. Bioresource Technology 289, 121744. 10.1016/j.biortech.2019.121744.

63. Brányiková, I., Maršálková, B., Doucha, J., Brányik, T., Bišová, K., Zachleder, V., and Vítová, M. (2011). Microalgae—novel highly efficient starch producers. Biotechnology and Bioengineering 108, 766–776. 10.1002/bit.23016.

64. Zhi, M., Zhao, Y., Zeng, X., Maddela, N.R., Xiao, Y., Chen, Y., Prasad, R., and Zhou, Z. (2023). Filamentous cyanobacteria and hydrophobic protein in extracellular polymeric substances facilitate algae—bacteria aggregation during partial nitrification. International Journal of Biological Macromolecules 251, 126379. 10.1016/j.ijbiomac.2023.126379.

65. Wang, Q.-R., Hong, Y., and Li, L.-H. (2023). Insights into differences between spore-assisted and pellet-assisted microalgae harvesting using a highly efficient fungus: Efficiency, high-value substances, and mechanisms. Science of The Total Environment 877, 162945. 10.1016/j.scitotenv.2023.162945.

66. Pertea, M., Pertea, G.M., Antonescu, C.M., Chang, T.-C., Mendell, J.T., and Salzberg, S.L. (2015). StringTie enables improved reconstruction of a transcriptome from RNA-seq reads. Nat Biotechnol 33, 290–295. 10.1038/nbt.3122.

