## supplementary_materials for "A shift between r and K strategies in the life cycle of a unicellular microalga"

For papers with only two authors: Paste the full name of the first and second author.
Lu Liu *et al.*

*Zhonghua Cai.

**This PDF file includes:**

Supplementary Text
Figs. S1 to S6

**Fig. S1.**

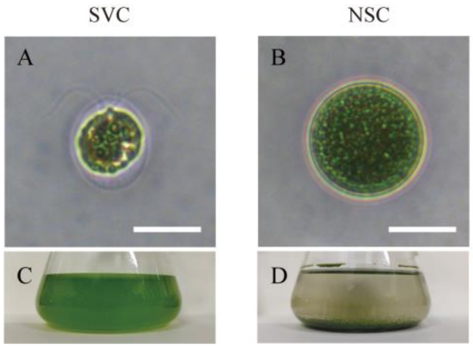


**Supplementary Figure 1. Images of SVC and NSC.** (**A**) The microscopic image of a SVC. (**B**) The microscopic image of an NSC. c The image of enriched SVC. d The image of enriched NSC. Scale bar: 20 μm.

**Fig. S2.**

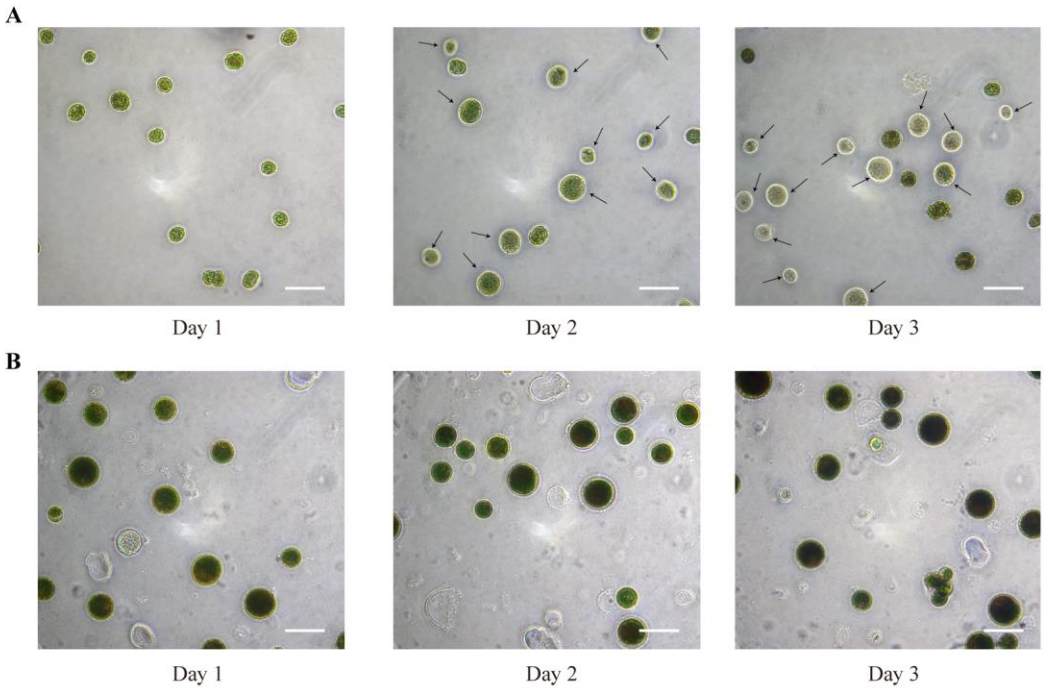


**Supplementary Figure 2. Cellular changes exposed high light irradiation.** (**A**) The performance of SVC exposed to high light irradiation. (**B**) The performance of NSC exposed to high light irradiation. The photobleached cells (dead cell) were indicated by the arrow. Scale bar: 50 µm.

**Fig. S3.**

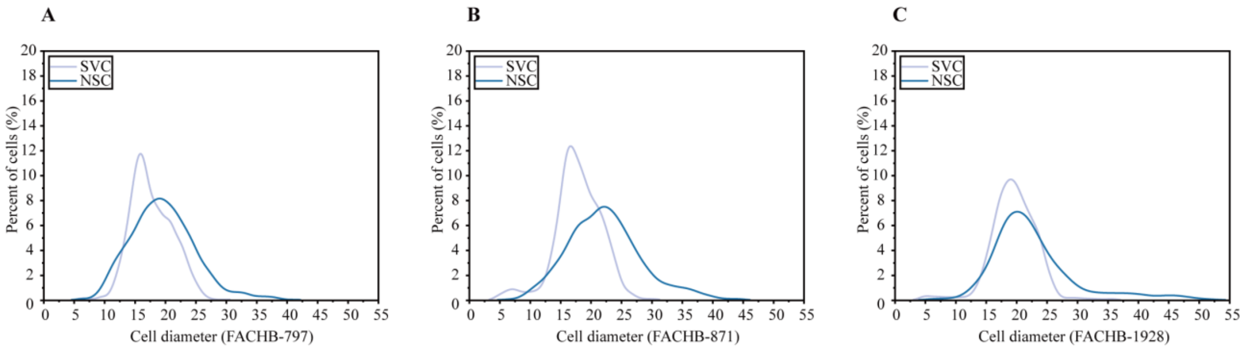


**Supplementary Figure 3. Cellular size distribution of two phenotypes of three other *H. lacustris strains*.** (**A**) The Gaussian kernel density curve of H. lacustris FACHB-797 (Hubei province, China). (**B**) The Gaussian kernel density curve of H. lacustris FACHB-871 (Jiangxi province, China). (**C**) The Gaussian kernel density curve of H. lacustris FACHB-1928 (Yunnan province, China). The light blue and dark blue colour represent SVC and NSC, respectively. For each phenotype, at least 1000 cells were measured.

**Fig. S4.**

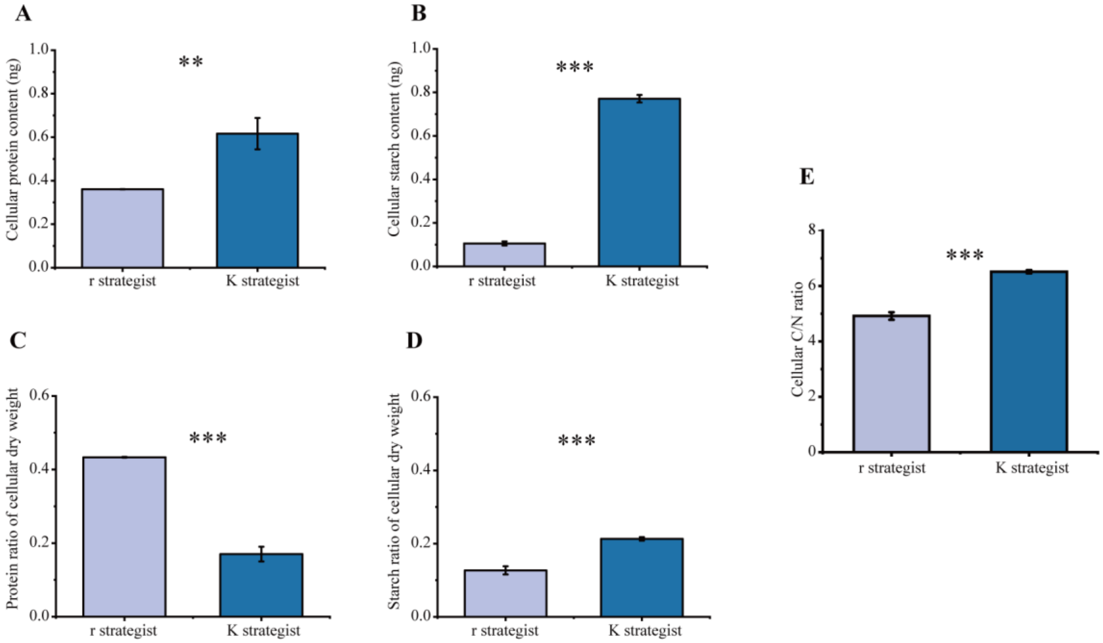


**Supplementary Figure 4. Physiological profiles of each strategist.** (**A**) The cellular protein content of each strategist. (**B**) The cellular starch content of each strategist. (**C**) The ratio of cellular protein to dry weight of each strategist. (**D**) The ratio of cellular starch to dry weight of each strategist. (**E**) Elemental carbon to nitrogen ratio of each strategist. The light blue and dark blue colour represent r strategists and K strategists, respectively. The error bars indicate the standard deviation of the triplicate values. N = 3 independent biological replicates. Statistical significance was calculated by Welch's *t* test for pairwise comparisons of two treatments. Significance: * ($p < 0.05$), ** ($p < 0.01$), *** ($p < 0.001$).

**Fig. S5.**

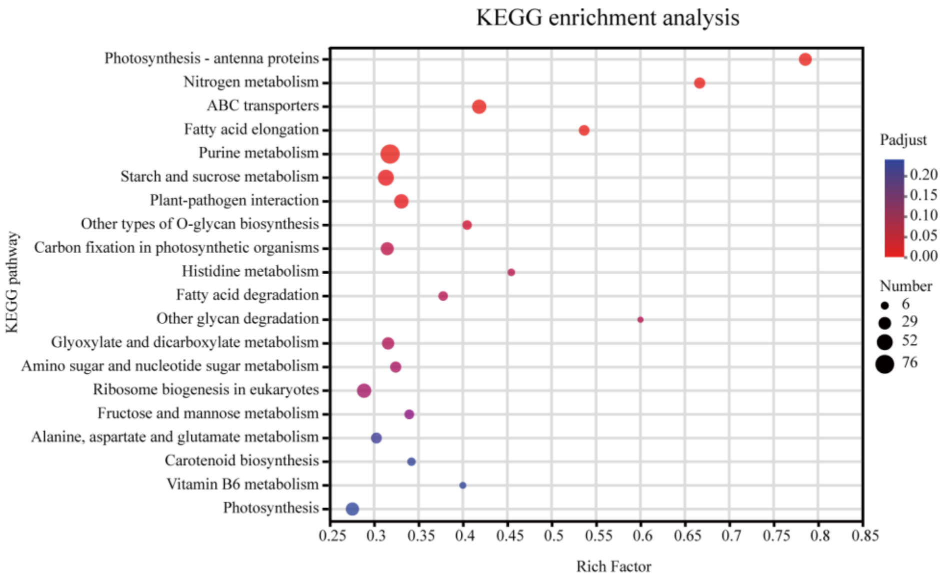


**Supplementary Figure 5. KEGG enrichment analysis.** Top 20 pathways were shown. DEGs with *p* value < 0.05 (right-tailed Fisher's exact test followed by a Benjamin-Hochberg adjustment), was used for enrichment analysis. The dot size indicates the number of genes for each pathway, and the colour bar indicates the *p* value. N = 3 independent biological replicates.

**Fig. S6.**

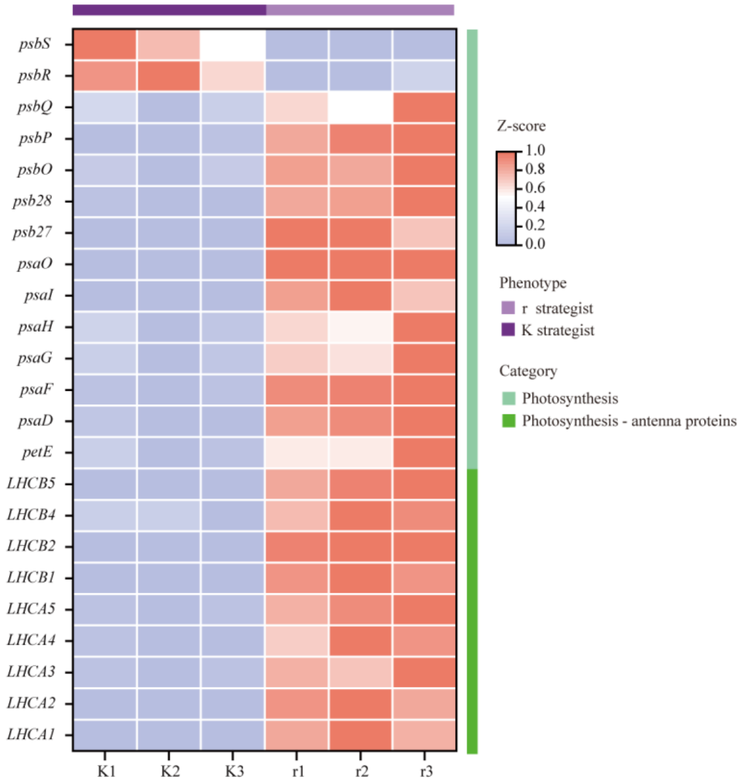


**Supplementary Figure 6. Heatmap analysis of DEGs involved in photosynthesis-antenna protein and photosynthesis.** The heatmap data is normalized, the blue colour represents downregulation, the red colour represents upregulation. The light purple bar represents r strategists (r), while the dark purple bar represents K strategists (K). Gene names are showed as KO names. Differentially expressed genes (DEGs) were identified by using DESeq2 with *q* value $\leqslant 0.05$, accompanied by an absolute value of $log_2FC$ ($log_2$ fold change) $\geqslant 1$. N = 3 independent biological replicates.

**Data S1. Raw data used in this manuscript (separate file)**
